# Defining and Evaluating Cell–Cell Relation Extraction from Biomedical Literature under Realistic Annotation Constraints

**DOI:** 10.64898/2025.12.01.691726

**Authors:** Mei Yoshikawa, Tadahaya Miuzuno, Yohei Ohto, Hiromi Fujimoto, Hiroyuki Kusuhara

## Abstract

Extracting cell–cell relations from biomedical literature is essential for understanding intercellular communication in immunity, inflammation, and tissue biology. However, cell–cell relation extraction has not been established as a standalone biomedical relation extraction task, and no benchmark corpus or systematic evaluation framework currently exists. Fully manual corpus construction is costly and difficult to scale, limiting literature-based analyses of cell–cell communication. Here, we define a sentence-level cell–cell relation extraction task and construct complementary manually annotated corpora under realistic annotation constraints. To enable scalable annotation, rule-based literature mining is used solely as an annotation accelerator to identify candidate sentences, while all relation labels are assigned manually. In addition, an independently annotated PubMed corpus without rule-based filtering is constructed to evaluate robustness on natural sentence distributions. Using these resources, we evaluate representative model configurations involving entity indication strategies, classification architectures, and continued pre-training. Our results show that cell–cell relation extraction remains challenging under realistic conditions. Increasing training data size yields consistent performance gains, and specific combinations of entity-aware architectures and continued pre-training provide modest robustness improvements. Nevertheless, performance on unfiltered PubMed sentences remains in the 70% accuracy range, and error analyses indicate that failures cannot be readily explained by simple surface-level factors. Comparisons with general-purpose large language models further suggest that task complexity, rather than model class, is the primary limiting factor. Together, these findings establish a practical foundation for literature-scale cell–cell relation extraction while clarifying its intrinsic limitations.

## 1 Introduction

Cell–cell communication (CCC) plays a central role in immunity, inflammation, tissue homeostasis, and disease progression^1^. Experimental and computational approaches, including single-cell and spatial transcriptomics, have enabled systematic analysis of intercellular interactions at the molecular level^2–4^. However, interpretation of such data still relies heavily on prior biological knowledge describing how specific cell types interact, regulate, or influence one another. Much of this knowledge is documented in the biomedical literature, making automated extraction of CCC-related information from text an important but largely unmet need^5–9^.

In biomedical literature, evidence relevant to CCC is often expressed as explicit or implicit relations between cell types. In this study, we refer to such textual descriptions as cell–cell relations and formulate their extraction from text as a biomedical relation extraction (BioRE) task. While BioRE has been extensively studied for relations involving well-defined entities such as proteins, chemicals, and diseases^10,11^, cell–cell relation extraction has not been established as a standalone task, and no benchmark corpus or evaluation framework currently exists^12^.

Several factors have hindered the formulation of cell–cell relation extraction. Cell types are described using heterogeneous and context-dependent nomenclature, often exhibit hierarchical organization, and frequently co-occur within the same sentence without implying a direct biological relationship^13,14^. These characteristics complicate both annotation and modeling, making direct transfer of existing BioRE datasets and methods non-trivial. As a result, literature-based approaches to CCC have not been systematically evaluated as an independent relation extraction problem despite their biological relevance. Beyond these conceptual challenges, constructing annotated data for CCC presents substantial practical difficulties. Identifying sentences that potentially describe cell–cell relations requires searching a vast literature, while assigning relation labels demands careful biological interpretation. For CCC, dataset construction therefore benefits from mechanisms that narrow down candidate sentences for manual review, while still allowing explicit labeling of both relation-positive sentences and sentences in which two cell types are mentioned without any relationship described.

Against this background, we aim to enable literature-scale CCC extraction under realistic annotation constraints. We define a cell–cell relation extraction task and establish annotation guidelines to distinguish biologically meaningful relations from simple co-occurrence. Using these guidelines, we construct two complementary manually annotated corpora: a SemMed^15^-based corpus, in which candidate sentences are identified by rule-based filtering and labeled manually, and an independently annotated PubMed corpus created without such filtering to evaluate robustness on natural sentence distributions. Using these resources, we evaluate representative sentence-level BioRE configurations involving different entity representations, classification architectures, and continued pre-training strategies. Rather than proposing a new state-of-the-art model or deriving general design principles for BioRE, our goal is to clarify what can currently be achieved for extracting literature-level evidence of CCC and to characterize the practical limits of sentence-level approaches under realistic data conditions.

Figure 1 provides an overview of the overall study design, including dataset construction, model configurations, and evaluation settings. To make large-scale manual annotation feasible under realistic constraints, we intentionally adopt rule-based literature mining not as a source of labels, but solely as an annotation accelerator to narrow down candidate sentences for expert review. All relation labels are assigned manually, and an independently annotated PubMed corpus without such filtering is used to explicitly assess robustness beyond the filtered setting.

**Figure 1.**
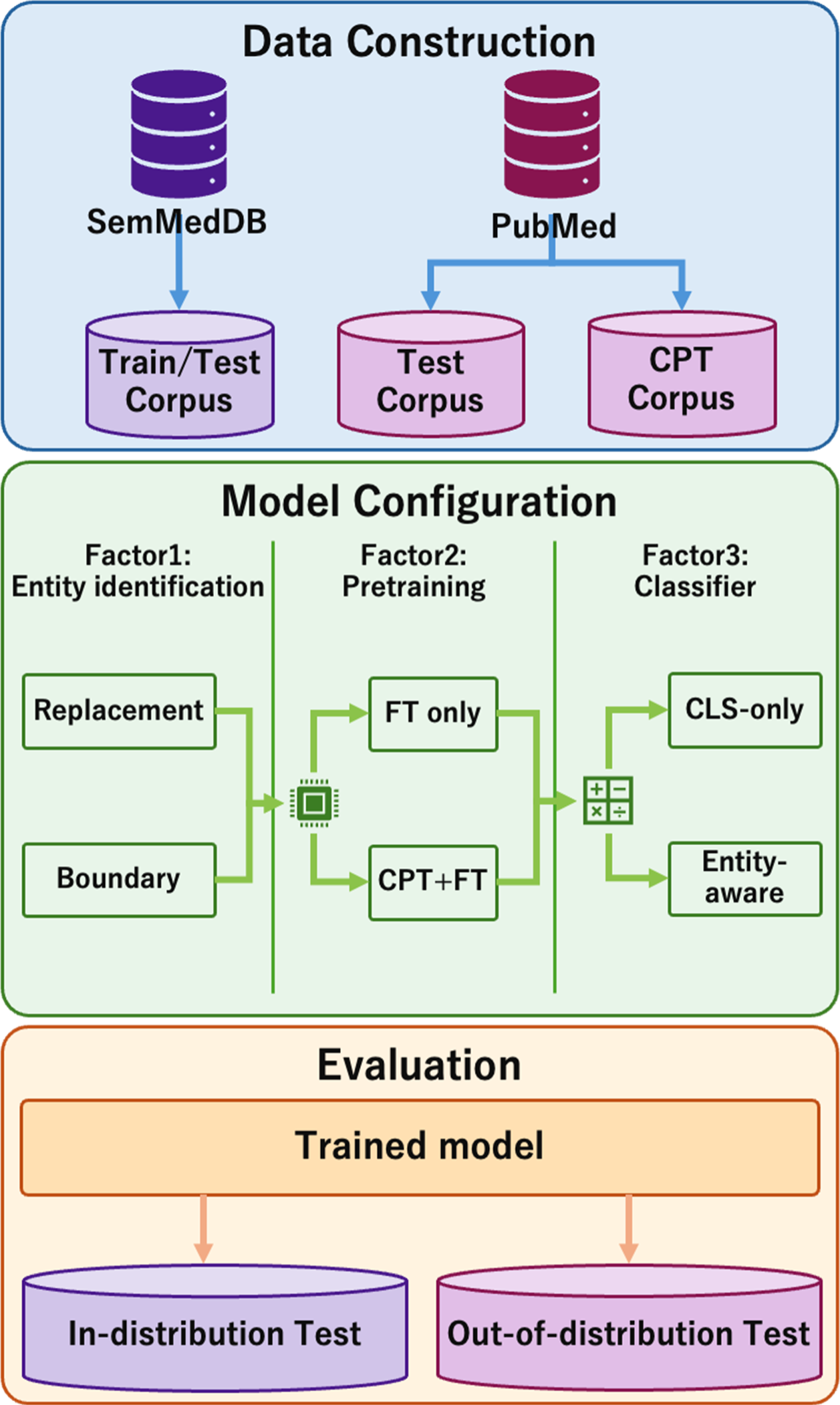
Schematic overview of the study design. The experimental workflow consists of three phases: (A) Dataset Construction, where purpose-specific datasets are constructed from SemMedDB and PubMed; (B) Model Configuration, systematically comparing combinations of Entity Indication (Replacement vs. Boundary), Pre-training Strategy (with/without CPT), and Classification Architecture; and (C) Evaluation, assessing performance on both in-distribution (SemMed-derived) and out-of-distribution (PubMed-derived) datasets.

## 2 Related Work

### 2.1 Entity marking and replacement strategies in BioRE

How target entities are specified in the input text is a fundamental design choice in relation extraction. Boundary marking approaches, such as those introduced in R-BERT^16^, explicitly denote entity spans and allow models to incorporate span-level representations^17–21^. In contrast, entity replacement strategies, which substitute entity names with category labels (e.g., CELL), have been widely adopted in biomedical NLP to reduce vocabulary variability caused by synonymy and terminological diversity.

Both strategies have been reported to be effective in different settings, but systematic comparisons remain limited in the biomedical domain. In particular, little is known about how these strategies behave for relations involving homogeneous entity types, such as cell–cell relations, where surface-level cues and co-occurrence patterns differ substantially from more commonly studied BioRE tasks.

### 2.2 CLS-only vs. entity-aware classification

BERT-based relation classification models typically fall into two categories: CLS-only architectures that rely on the global [CLS] representation, and entity-aware architectures that explicitly incorporate representations of target entity spans in addition to the [CLS] token. CLS-only models offer simplicity and have been shown to provide strong baselines in various relation extraction tasks^22,23^, while entity-aware approaches have been reported to better capture local information around entity mentions^24^.

However, the relative effectiveness of CLS-only and entity-aware architectures has not been systematically established across relation extraction tasks, particularly in biomedical text where sentence structures are complex and multiple entity mentions often appear. For cell–cell relation extraction, which involves relations between identical entity types, no comparative evaluation has previously been reported, leaving this design choice largely unexplored in this context.

### 2.3 Special tokens and continued pre-training

Entity-aware models often rely on special boundary markers that are not present in the original pre-training vocabulary of BERT-based encoders^25^. This mismatch raises the question of whether fine-tuning alone is sufficient to learn meaningful representations for such markers, or whether additional pre-training on marker-containing text is beneficial. Continued pretraining (CPT) has been reported to improve performance in some general-domain and biomedical NLP tasks by adapting models to task-specific input formats^21,26–30^.

Despite these reports, the effect of CPT in combination with domain-specific encoders such as BiomedBERT, particularly for boundary-marked BioRE tasks, has not been systematically examined. Its interaction with different classification architectures in cell–cell relation extraction therefore remains an open question.

### 2.4 Cell–cell relation extraction from literature

BioRE has been extensively studied for interactions such as protein–protein, chemical–disease, and gene–protein relations, supported by established benchmarks and shared tasks^11,31^. In contrast, no dedicated task formulation or annotated corpus exists for extracting cell–cell relations directly from biomedical literature.

As a result, key methodological questions—such as how to represent cell entities in text and which model architectures are most suitable—have not been empirically evaluated for this class of relations. Addressing this gap requires both the construction of appropriate datasets and the evaluation of representative BioRE configurations tailored to cell–cell relations.

### 2.5 SemMedDB as a candidate sentence filtering resource

SemMedDB^15^ consists of PubMed sentences associated with subject–predicate–object tuples automatically extracted by the rule-based SemRep system^32^. Rather than serving as a source of labeled training data, SemMedDB provides a mechanism for identifying candidate sentences in which relations between biomedical entities are likely to be described.

Because SemRep preferentially extracts sentences with relatively explicit predicate structures, sentences derived from SemMedDB may differ in distribution from unfiltered PubMed text. However, the implications of using SemMedDB for candidate sentence selection—followed by full manual annotation—have not been systematically evaluated for cell–cell relation extraction, motivating a direct comparison with independently sampled PubMed sentences.

## 3 Datasets

Due to the absence of existing benchmarks for cell–cell relation extraction, we constructed datasets specifically for this study (**Supplementary Table 1**). These consist of two manually annotated corpora for model training and evaluation, and an additional unlabeled corpus used exclusively for CPT. The overall dataset construction pipeline is illustrated in Figure 2. Explicit annotation guidelines were defined in advance, and all relation labels in the annotated corpora were assigned manually.

**Figure 2.**
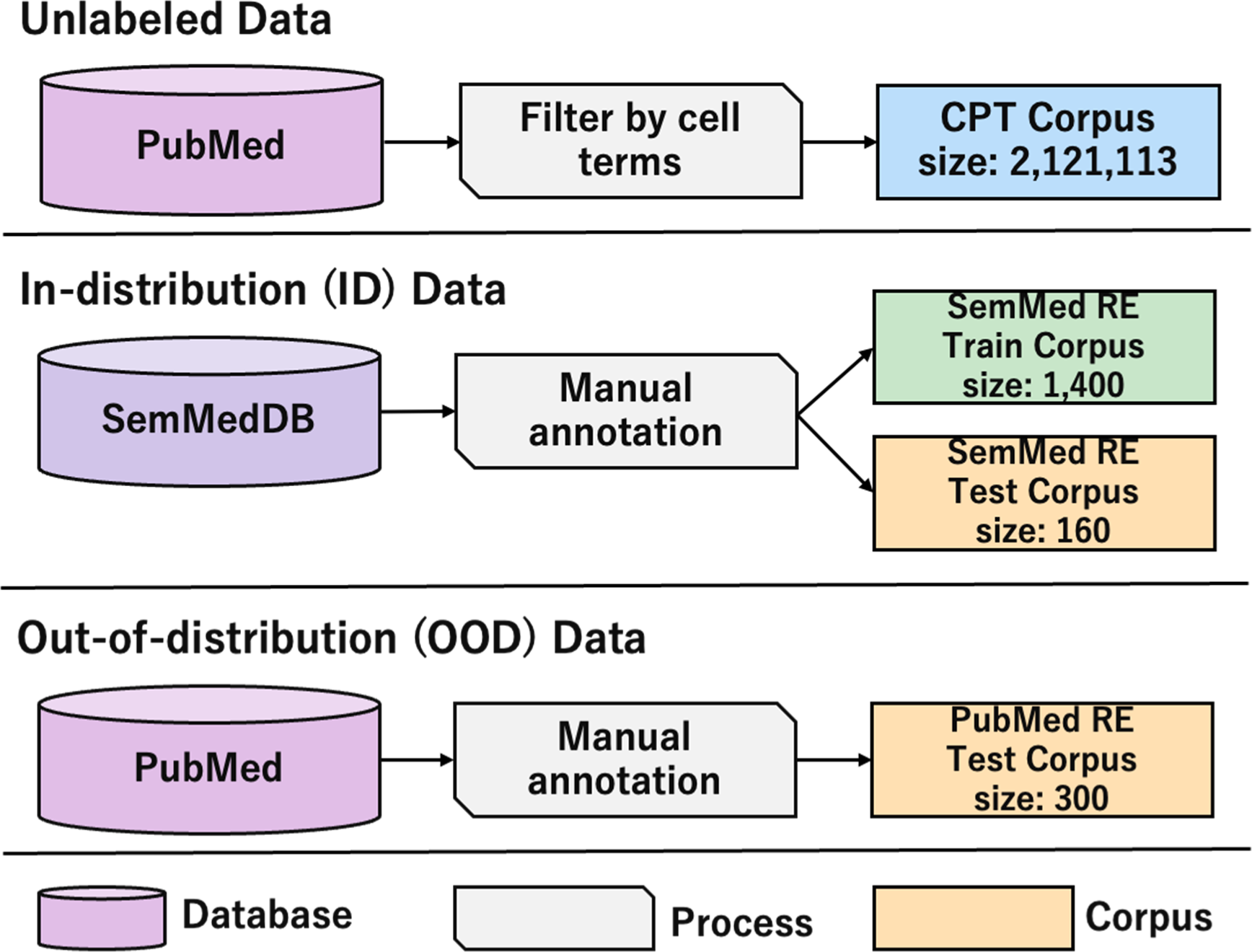
Dataset construction pipeline. The figure illustrates the process of generating final corpora from their respective data sources. Top: Construction of the unlabeled corpus for Continued Pre-training (CPT) via dictionary-based filtering from PubMed. Middle: Construction of the supervised training set and In-Distribution (ID) test set utilizing Predications (triples) from SemMedDB. Bottom: Construction of the Out-of-Distribution (OOD) test set via dictionary-based matching and random sampling from PubMed. “Size” denotes the number of sentences included in each final dataset.

**Figure 3.**
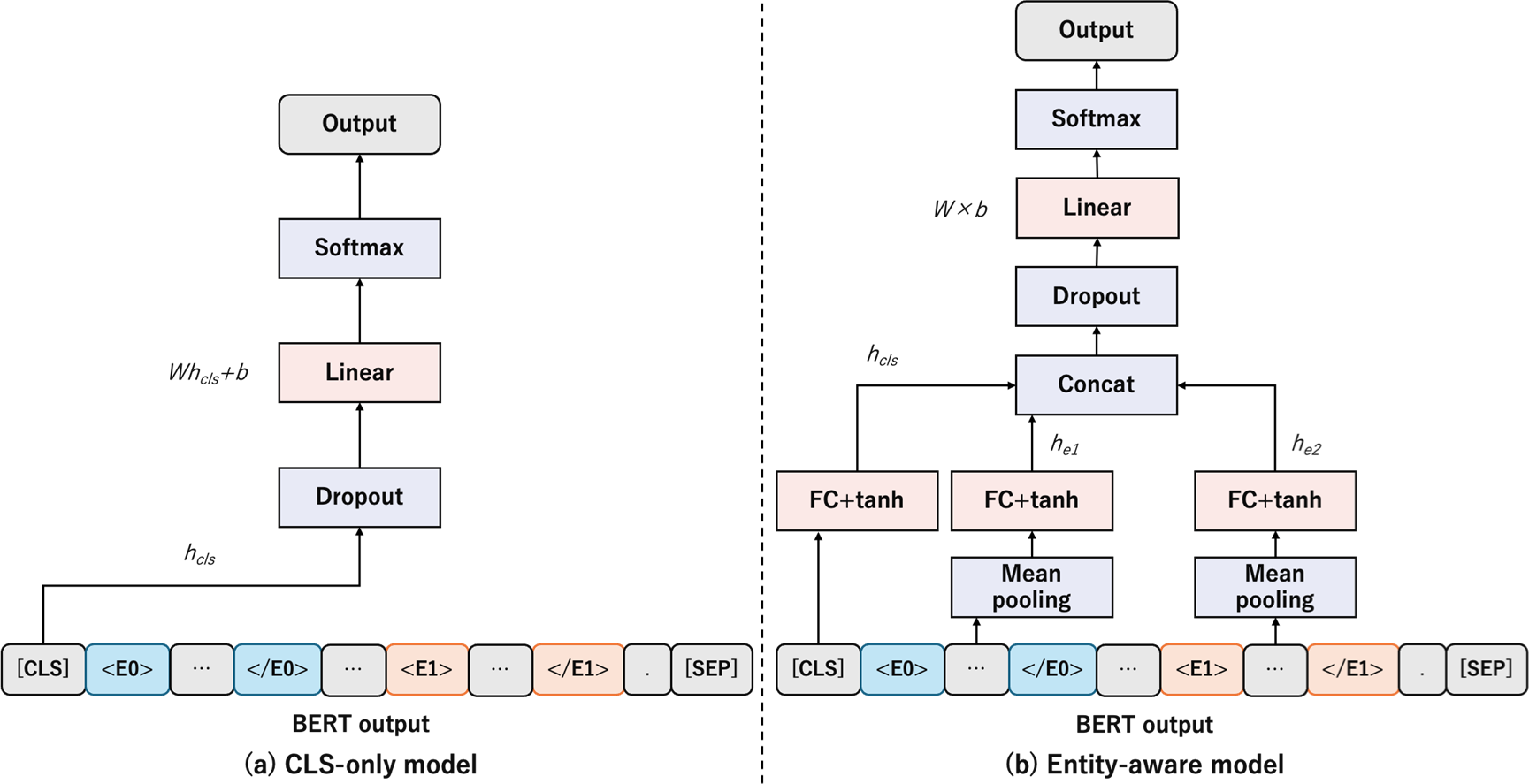
Schematic of classification architectures. (a) CLS-only model: Performs classification utilizing only the global representation corresponding to the [CLS] token. (b) Entity-aware model: In addition to the [CLS] token, entity-specific representations are extracted via mean pooling of token embeddings within the entity spans (or special tokens in the Replacement method). These vectors are concatenated and passed through a fully connected (FC) layer and a softmax function for binary classification.

### 3.1 Annotation guideline and process

Definitions of cell–cell interactions vary across literature, necessitating criteria to distinguish substantial relationships from simple co-occurrences. In this study, a relationship was defined as a description explicitly indicating biological influence between two cells and was classified into three types:

1. Unidirectional interaction: Interaction directed from one cell to another.
2. Bidirectional interaction: Interactions are described bidirectionally.
3. Change/transition: Descriptions of cell state changes, such as conversion, differentiation, or transition.

The guidelines included definitions, decision criteria, and both typical and atypical examples. Consistency was ensured through an iterative process of drafting, annotating, reconciling rules, and revising the guidelines. The final annotation was performed based on these established guidelines (refer to **Supplementary Information** for details).

### 3.2 SemMed-filtered annotated corpus for training and in-distribution evaluation

The annotated corpus for model training and in-distribution evaluation was constructed using the SemMedDB^15^. In this study, SemMedDB was used solely as a candidate sentence filtering resource to identify sentences in which two cell types are mentioned in contexts suggestive of relations.

All candidate sentences were subsequently annotated manually according to the guidelines described in **Supplementary Information**. Both positive instances, where a biologically meaningful cell–cell relation is described, and negative instances, where two cell types are mentioned without an indicated relation, were labeled explicitly. Using this procedure, we obtained a manually annotated corpus of 1,560 sentences, which was split into 1,400 sentences for training and development and 160 sentences for in-distribution (ID) testing. **Supplementary Figure 1** shows the distribution of the above three types in this corpus.

### 3.3 PubMed-based natural text corpus for test

To evaluate model robustness beyond the SemMed-based setting, we constructed an independent test corpus from PubMed abstracts without SemMedDB or SemRep filtering. PubMed abstracts were segmented into sentences, and sentences containing at least two cell-type mentions were identified using a cell-type dictionary. From this pool, 680 sentences were randomly sampled and manually annotated in advance using fixed random seeds.

All sentences in this corpus were annotated manually using the same guidelines as in **Supplementary Information**. Because this corpus is derived without rule-based filtering, it reflects the linguistic and structural diversity encountered in practical literature mining and serves as an out-of-distribution evaluation set. Note that although all 680 sentences were fully annotated, model evaluation was performed by randomly sampling 300 sentences per run from this annotated pool, using fixed random seeds for reproducibility. This design allows robustness to be assessed across different subsets of naturally occurring PubMed sentences, rather than relying on a single fixed test set.

### 3.4 PubMed-based unlabeled corpus for continued pretraining

For CPT, we constructed an unlabeled corpus from PubMed abstracts. Approximately two million sentences containing cell-type mentions were extracted using the same dictionary as in Section 3.3. For each sentence, two cell mentions were selected, and entity markers were inserted to match the input format used by entity-aware models.

This corpus was used to expose the encoder to marker-containing sentence structures prior to fine-tuning, thereby adapting the model to the downstream input format. The CPT procedure followed standard masked language modeling, with marker tokens excluded from masking.

## 4 Methods

In this study, we examine representative modeling choices that are required to instantiate a sentence-level cell–cell relation extraction task, focusing on how different design decisions affect performance under realistic data conditions (Figure 1). Specifically, we consider entity indication methods, classification architectures, and the use of CPT for handling special tokens.

### 4.1 Problem formulation

The input consists of a sentence *x* extracted from scientific literature and two cell-type entities, *e*_l_ and *e*_2_, appearing within it. The output is a binary label *y* ∈ {0, 1} indicating whether a cell–cell interaction is described between *e*_l_ and *e*_2_. In cell–cell relation extraction, both entities belong to the same semantic category, and the relationship can be symmetric. Moreover, cell names exhibit high lexical diversity and strong context dependence, and simple co-occurrence of two cell types does not necessarily imply a meaningful interaction. As a result, accurate prediction requires careful interpretation of the surrounding textual context.

These characteristics directly influence the design of relation extraction models, including choices related to entity indication, representation extraction, and pre-training strategies.

### 4.2 Entity marking, model architecture, and pretraining strategies

Multiple models were constructed by combining entity indication methods, classification architectures, and the presence or absence of CPT.

#### Replacement (Category substitution)

In this method, cell names in the sentence are replaced with category labels such as [CELL0] and [CELL1] (**Supplementary Figure 2a**).

Macrophages activate T cells. → [CELL0] activate [CELL1].

This approach reduces vocabulary variance and mitigates the out-of-vocabulary (OOV) problem. It may discard fine-grained biological semantics, but can improve robustness under lexical variability.

#### Boundary marking (R-BERT style)

Target entities are enclosed by markers, such as <E0>…</E0> and <E1>…</E1> (**Supplementary Figure 2b**).

Macrophages activate T cells. → <E0>Macrophages</E0> activate <E1>T cells</E1>.

This allows the model to identify explicit entity spans and extract representations from the surrounding context. A challenge, however, is that these special tokens are absent during the original pre-training of BiomedBERT; thus, the learning of their embeddings depends solely on fine-tuning.

#### CLS-based classifier

This architecture inputs the global [CLS] representation into the classifier (Figure 2a).

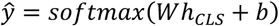

While the structure is simple and training is stable, it does not explicitly utilize local entity information, which may limit the model’s ability to explicitly leverage local information around target entities in context-dependent tasks such as cell–cell RE.

#### Entity-aware classifier (R-BERT)

This architecture incorporates entity-specific representations in addition to the global [CLS] token (Figure 2b). The extraction method depends on the entity indication strategy:

When boundary marking is utilized, entity representations are extracted via average pooling of the tokens enclosed by the markers:

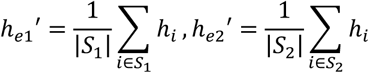

When replacement is utilized, the final hidden states corresponding to the special tokens [CELL0] and [CELL1] are directly used as ℎ_*e*l_′ and ℎ_*e*2_′.

Next, these averaged representations are passed through a separate fully connected layer with weights *W_e_* and bias *b_e_*, followed by a tanh activation to obtain the final entity representations:

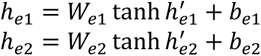

The final prediction is obtained by concatenating the [CLS] vector and the two span vectors, followed by input into an MLP.

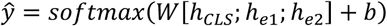

The advantage of this approach is the ability to directly handle local information specific to the target entities.

#### Continued Pretraining

The motivation for CPT was to expose the encoder to input formats containing entity markers prior to fine-tuning, thereby adapting the model to the structural characteristics of the downstream task. CPT was evaluated for both CLS-only and entity-aware architectures to assess whether such adaptation affects performance in cell–cell relation extraction (**Supplementary Figure 3**).

### 4.3 Fine-tuning

For fine-tuning, the cell–cell RE corpus of 1,400 sentences constructed from SemMed Predications was used. Implementation was carried out using the Hugging Face Transformers library^33^. The parameters of BiomedBERT (base or CPT version) were updated to optimize for the relation classification task.

Optimization was performed using the RAdamScheduleFree^34^ optimizer with a batch size of 16 for fine-tuning and 128 for CPT, and a learning rate of 2 × 10⁻⁵ for both phases. Training was run for up to 20 epochs during fine-tuning and up to 2 epochs during CPT, with early stopping applied based on development set loss. To ensure reproducibility, all experiments were conducted with multiple random seeds.

### 4.4 Evaluation

The test corpus of 680 cell–cell RE sentences extracted and annotated from PubMed was used as the source for evaluation. Since this dataset is derived from a natural text distribution different from SemMed, it allows for the measurement of out-of-distribution (OOD) generalization performance.

In addition to BERT-based models, we evaluated several general-purpose large language models (LLMs) on the same OOD PubMed test set to provide a contextual reference for task performance. LLMs were prompted to perform sentence-level binary classification without any task-specific fine-tuning or additional training (**Supplementary Figure 4**). This comparison is intended to contextualize the performance of task-specific models rather than to establish direct competitiveness with LLMs optimized for different objectives.

Performance was measured using accuracy, F1 score, and Matthews correlation coefficient (MCC). Among these, MCC was used as the primary evaluation metric due to its robustness to class imbalance^35^.

### 4.5 Model configurations

The models compared in this study consist of 8 conditions derived from the combination of three axes: Entity Indication, Classification Architecture, CPT (**Supplementary Table 2**). This configuration allows us to examine the effects of each design element and their interactions within the scope of cell–cell relation extraction.

## 5 Results

### 5.1 In-distribution and out-of-distribution performance

We first compared model performance under ID and OOD evaluation settings to examine how design choices manifest under different data selection regimes. The ID test set was derived from SemMedDB^15^, where candidate sentences were pre-selected by the rule-based SemRep system^32^. The OOD test set, in contrast, was sampled directly from PubMed using dictionary-based matching without predicate-based filtering. Importantly, all labels in both settings were manually annotated using identical guidelines, ensuring comparable ground truth quality.

Across all configurations, models achieved consistently high performance on the ID test set, with accuracy values clustered around 0.82 (**Table 1**). Although modest differences among configurations were observed, the overall performance range under ID evaluation was narrow. On the OOD PubMed test set, performance decreased across all configurations, with accuracy values spanning a broader range, approximately from 0.70 to 0.76 (**Table 2**).

**Table 1.** Performance under in-distribution (ID) evaluation on the SemMed dataset. The table reports Accuracy, Matthews Correlation Coefficient (MCC), and F1-score for all model configurations evaluated on the in-distribution (ID) SemMed test set. Results are presented as mean ± standard deviation across multiple runs.

| Model | Accuracy | MCC | F1 |
| --- | --- | --- | --- |
| R-ENT-CPT | **0.843±0.041** | **0.690±0.081** | **0.852±0.037** |
| R-ENT-base | 0.840±0.053 | 0.682±0.106 | 0.845±0.052 |
| R-CLS-CPT | 0.816±0.030 | 0.637±0.059 | 0.825±0.028 |
| R-CLS-base | 0.818±0.030 | 0.638±0.059 | 0.826±0.028 |
| B-ENT-CPT | 0.825±0.057 | 0.655±0.115 | 0.836±0.053 |
| B-ENT-base | 0.820±0.034 | 0.642±0.068 | 0.825±0.036 |
| B-CLS-CPT | 0.829±0.038 | 0.662±0.076 | 0.839±0.035 |
| B-CLS-base | 0.840±0.023 | 0.684±0.046 | 0.848±0.021 |

**Table 2.** Performance under out-of-distribution (OOD) evaluation on the PubMed dataset. The table reports Accuracy, Matthews Correlation Coefficient (MCC), and F1-score for all model configurations evaluated on the out-of-distribution (OOD) PubMed test set. Results are presented as mean ± standard deviation across multiple runs.

| Model | Accuracy | MCC | F1 |
| --- | --- | --- | --- |
| **R-ENT-CPT** | **0.757±0.009** | **0.523±0.020** | **0.734±0.009** |
| R-ENT-base | 0.731±0.015 | 0.474±0.032 | 0.699±0.017 |
| R-CLS-CPT | 0.715±0.021 | 0.435±0.044 | 0.695±0.018 |
| R-CLS-base | 0.713±0.025 | 0.431±0.053 | 0.694±0.019 |
| B-ENT-CPT | 0.745±0.012 | 0.498±0.024 | 0.719±0.015 |
| B-ENT-base | 0.703±0.008 | 0.415±0.017 | 0.668±0.016 |
| B-CLS-CPT | 0.738±0.023 | 0.480±0.051 | 0.722±0.017 |
| B-CLS-base | 0.743±0.011 | 0.492±0.023 | 0.724±0.013 |

Comparing ID and OOD evaluations reveals a systematic performance degradation when models are applied to data drawn from a different source, with ID performance consistently exceeding OOD performance across configurations (**Table 3**). In addition, while differences among configurations are present under both settings, the range of mean performance values is wider under OOD evaluation than under ID evaluation. This observation suggests that the OOD setting provides a less constrained evaluation regime, in which variations in model behavior become more apparent than in the highly structured, rule-based filtered ID setting.

**Table 3.** Generalization gap between in-distribution (ID) and out-of-distribution (OOD) evaluations. The table summarizes the performance gap between ID and OOD evaluations for each model configuration. Δ represents the generalization gap, calculated as Score_SemMed_−Score_PubMed_. Variance estimates are not reported, as the gap is used as a descriptive indicator of robustness rather than as a statistical estimator.

| Model | ΔAccuracy | ΔMCC | ΔF1 |
| --- | --- | --- | --- |
| R-ENT-CPT | 0.085 | 0.167 | 0.118 |
| R-ENT-base | 0.109 | 0.208 | 0.145 |
| R-CLS-CPT | 0.101 | 0.202 | 0.130 |
| R-CLS-base | 0.104 | 0.207 | 0.131 |
| B-ENT-CPT | 0.080 | 0.157 | 0.117 |
| B-ENT-base | 0.117 | 0.227 | 0.157 |
| B-CLS-CPT | 0.091 | 0.182 | 0.116 |
| B-CLS-base | 0.097 | 0.192 | 0.124 |

### 5.2 Interaction between classification architecture and continued pre-training

We next examined the interaction between classification architecture and CPT, explicitly contrasting their behavior under ID and OOD evaluation. Under ID evaluation, differences attributable to CPT or architecture were limited (**Table 1**). In both CLS-only and entity-aware architectures, models trained with and without CPT showed largely overlapping performance distributions, and the observed differences were small relative to the corresponding standard deviations. Consequently, the ID setting did not allow for a clear separation of model configurations based on CPT or architectural choice.

Under OOD evaluation, clearer differences emerged (**Table 2**). For CLS-only architectures, the introduction of CPT resulted in minimal changes in performance. In contrast, entity-aware architectures consistently showed improved performance when combined with CPT. In the Replacement setting, CPT increased OOD accuracy from 0.731±0.015 to 0.757±0.009, while in the Boundary setting, accuracy increased from 0.703±0.008 to 0.745±0.012. Similar trends were observed for MCC and F1.

These results indicate that the effect of CPT depends on the classification architecture. Rather than providing uniform improvements, CPT primarily benefits architectures that explicitly incorporate entity-level representations, and this interaction becomes apparent under OOD evaluation.

### 5.3 Interaction between entity indication strategy and continued pre-training

We next examined the effect of entity indication strategies, focusing on their interaction with CPT across evaluation settings. Under ID evaluation, differences between Replacement and Boundary marking were small regardless of whether CPT was applied, and performance distributions largely overlapped (**Table 1**). As a result, the effect of entity indication strategy was not clearly distinguishable under the source-matched ID setting.

In contrast, clearer differences emerged under OOD evaluation, particularly in entity-aware architectures (**Table 2**). When CPT was not applied, Replacement achieved substantially higher OOD accuracy than Boundary marking (0.731±0.015 vs. 0.703±0.008), with non-overlapping standard deviations. This result indicates that, in the absence of CPT, Boundary marking is more sensitive to source-shifted evaluation than Replacement. When CPT was applied, the performance gap between the two entity indication strategies was reduced. With CPT, Replacement achieved an OOD accuracy of 0.757±0.009 compared to 0.745±0.012 for Boundary marking, and the corresponding performance distributions partially overlapped. This suggests that CPT mitigates, but does not fully eliminate, the sensitivity of Boundary marking under OOD evaluation.

Taken together, these results indicate that the effect of entity indication strategy cannot be interpreted independently of pre-training strategy. Rather, CPT plays a compensatory role for Boundary marking, reducing the performance gap relative to Replacement when models are evaluated under source-shifted conditions.

### 5.4 Generalization gap analysis

To summarize the impact of source shift across evaluation settings, we report the generalization gap, defined as the difference between ID and OOD performance for each model configuration (**Table 3**). Across all configurations, performance decreased when models were evaluated on the OOD test set, resulting in a non-zero generalization gap for all metrics. This observation indicates that models trained on SemMed-filtered annotated data do not fully retain their performance when applied to PubMed sentences sampled without rule-based filtering.

Because ID and OOD evaluations were conducted with the same number of runs (N=5), the uncertainty of the generalization gap can be estimated by standard error propagation. Under this estimation, the uncertainty of the gap is comparable in magnitude to the gap itself for most configurations. Consequently, we do not perform statistical comparisons of gap magnitude across models, and instead treat the reported gaps as descriptive summaries of performance change between evaluation settings.

When interpreted alongside the OOD performance results in Sections 5.2 and 5.3, the gap estimates are consistent with the observation that CPT primarily benefits entity-aware architectures under OOD evaluation, whereas CLS-only architectures show limited sensitivity to CPT. At the same time, substantial gaps remain across all configurations, indicating that architectural and pre-training choices alone are insufficient to fully address the challenges posed by source-shifted evaluation in sentence-level cell–cell relation extraction.

To further examine whether the observed performance limitations could be attributed to simple surface-level factors, we conducted additional analyses stratified by training data size, sentence length, entity distance, and the presence of negation or speculative expressions (**Supplementary Figures 5–8, Supplementary Tables 3–4**). As expected, increasing the size of the manually annotated training data resulted in consistent performance improvements. However, no clear or systematic performance stratification was observed with respect to sentence length, entity distance, or the presence of negation and speculative cues. These results suggest that CCC extraction performance cannot be readily explained by individual surface-level properties alone.

### 5.5 Comparison with large language models

Finally, we compared the best-performing BERT-based configuration—an entity-aware architecture with Replacement and CPT—with several general-purpose large language models (LLMs) evaluated on the same OOD test set (**Figure 4, Supplementary Table 5**). This comparison is intended to contextualize performance rather than to establish a direct benchmark, as the models differ substantially in scale, training objectives, and inference cost.

**Figure 4.**
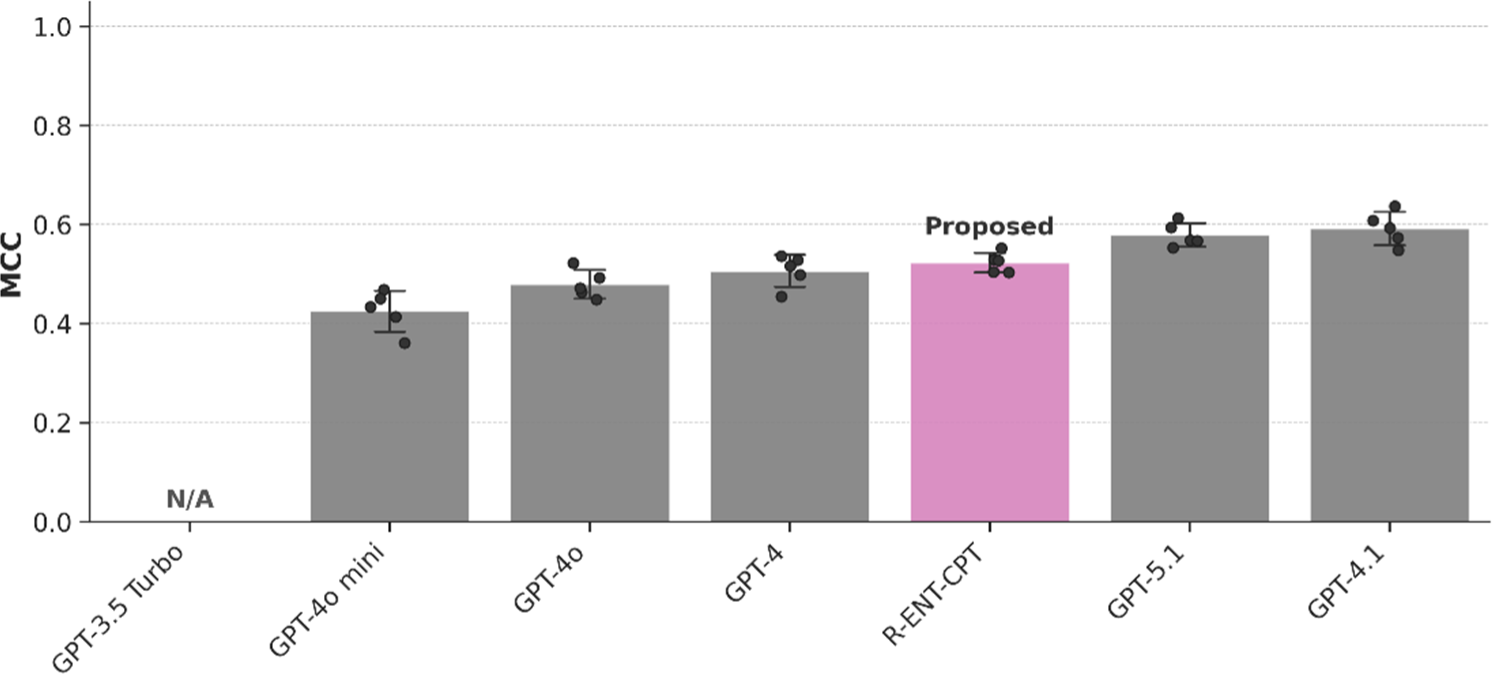
Performance comparison with Large Language Models (LLMs). The figure compares the proposed task-specific model (R-ENT-CPT) with several general-purpose LLMs on the out-of-distribution (OOD) PubMed test set using Matthews Correlation Coefficient (MCC). LLMs are evaluated in a prompt-based manner without task-specific fine-tuning and are included solely as reference baselines for contextual comparison. This analysis is not intended to claim superiority of the proposed model over LLMs, but rather to illustrate that sentence-level cell–cell relation extraction remains challenging even for large, general-purpose models under the same evaluation setting. Error bars indicate the standard deviation across multiple runs.

The proposed model achieved higher accuracy (0.757±0.009) than GPT-4o (0.715±0.012) and GPT-4o-mini (0.710±0.022), and performance comparable to GPT-4. More recent models, such as GPT-4.1 (0.795±0.016) and GPT-5.1 (0.789±0.012), achieved higher accuracy, but require substantially greater computational resources and rely on API-based access, which limits reproducibility and scalability in large-scale literature mining.

These results indicate that lightweight, domain-adapted discriminative models can remain competitive for structured information extraction from biomedical literature, particularly in large-scale settings where computational cost, scalability, and independence from external APIs are practical considerations.

## 6 Discussion

### 6.1 Enabling literature-scale cell–cell relation extraction under realistic constraints

The primary contribution of this study is to enable literature-scale cell–cell relation extraction under realistic annotation constraints. To our knowledge, no established task definition or annotated corpus has previously existed for extracting cell–cell relations directly from biomedical literature, despite their central importance in immunology, inflammation, and tissue biology.

By defining an explicit CCC task, constructing annotation guidelines, and assembling complementary annotated corpora, we demonstrate that CCC extraction is feasible in practice. Candidate sentences are identified using rule-based literature mining to reduce annotation cost, and all relations are annotated manually. Although overall performance remains limited, lightweight discriminative models and general-purpose large language models reach a similar performance range on this task, indicating that CCC extraction is non-trivial even for large models while remaining operationally tractable without reliance on API-based systems (**Figure 4, Supplementary Table 5**)^36^.

In practical terms, the present study establishes that literature-scale CCC extraction can be operationalized under realistic constraints, and that remaining performance limitations are driven more by intrinsic task complexity than by model class alone.

### 6.2 Rule-based candidate filtering as a practical annotation accelerator

A central methodological choice in this study was the use of SemMedDB^15^ as a source of candidate sentences for corpus construction. SemRep^32^, the rule-based system underlying SemMedDB, extracts predicate–argument structures using predefined linguistic and semantic patterns. While such rule-based extraction is necessarily limited and does not capture the full linguistic diversity of biomedical literature, it offers a practical mechanism for identifying candidate sentences and reducing annotation cost.

In the context of CCC, rule-based candidate filtering should be understood as an annotation accelerator rather than a labeling mechanism. Leveraging SemMedDB enabled efficient identification of candidate sentences, while all relations were annotated manually. In parallel, we constructed an independently annotated PubMed corpus without rule-based filtering to serve as a complementary evaluation benchmark. Although smaller in size, this corpus is comparable to those used in many existing BioRE benchmarks and is sufficient to reveal substantial performance differences between filtered and unfiltered sentence distributions.

### 6.3 What our results imply specifically for CCC

Our evaluation shows that common BioRE design choices exhibit conditional effects in the context of CCC. CPT improved robustness for entity-aware architectures under OOD evaluation, while its impact on CLS-only models was limited (**Table 1**). Similarly, the effect of entity indication strategies depended on the presence of CPT, with boundary-based representations showing increased sensitivity in its absence.

Importantly, these observations are specific to CCC and to the data construction regime examined in this study. They should not be interpreted as general design principles for biomedical relation extraction. Rather than identifying optimal architectures in a broad sense, the present results narrow the space of plausible design choices for CCC and clarify which directions are unlikely to yield substantial gains through sentence-level modeling alone.

### 6.4 Task-specific challenges in cell–cell relation extraction

Beyond methodological considerations, CCC presents intrinsic challenges that distinguish it from more established relation extraction settings. Cell types are often described using variable nomenclature and exhibit hierarchical organization. For example, a term such as “T cell” may subsume multiple subtypes, including CD4⁺ and CD8⁺ T cells, depending on context^13,14^. This variability in granularity and naming complicates sentence-level extraction, as models must implicitly resolve ontological ambiguity without explicit access to hierarchical knowledge.

The persistent performance limitations observed in this study suggest that further architectural refinement at the sentence level may yield diminishing returns for CCC extraction. Instead, explicitly incorporating cell-type hierarchies, resolving granularity mismatches, or leveraging document-level context and background biological knowledge may represent more promising directions for advancing CCC extraction from biomedical literature.

Consistent with this view, supplementary analyses indicate that CCC extraction errors are not strongly associated with simple surface-level factors such as sentence length, entity distance between cell mentions, or the presence of negation and speculative expressions (**Supplementary Figures 5–8, Supplementary Tables 3–4**). Even when the size of the manually annotated training data is increased, performance improvements remain incremental. Together, these observations reinforce the notion that the difficulty of CCC extraction arises from a combination of factors, including semantic variability and ontological ambiguity, rather than from isolated linguistic phenomena that can be addressed by straightforward modeling adjustments.

## 7 Conclusion

This study establishes a practical framework for extracting cell–cell relations from biomedical literature by defining the task, constructing complementary manually annotated corpora under realistic annotation constraints, and providing systematic baseline evaluations. Candidate sentences are identified using rule-based literature mining to reduce annotation cost, while all relation labels are assigned through manual annotation.

Rather than proposing a new state-of-the-art model, we clarify what can currently be achieved for CCC extraction using sentence-level BioRE approaches. Our results show that, despite incremental gains from specific design combinations, substantial performance limitations persist when models are evaluated on unfiltered PubMed sentences, indicating that CCC extraction remains intrinsically challenging.

By clarifying these limits and releasing task definitions, annotation guidelines, datasets, and baseline results, this work provides a concrete foundation for future studies on CCC extraction and related literature mining tasks involving complex and weakly defined biomedical entities.

## Author Contribution

Mei Yoshikawa: Methodology, Software, Investigation, Writing – Original Draft, Visualization.

Tadahaya Mizuno: Conceptualization, Resources, Supervision, Project administration, Writing – Original Draft, Writing – Review & Editing, Funding acquisition.

Yohei Ohto: Methodology, Software. Hiromi Fujimoto: Investigation Hiroyuki Kusuhara:Writing - Review

## Availability

Code and models are available at https://github.com/mizuno-group/cc-bert.

## Acknowledgement

We thank the contributors to the SemMedDB database utilized in this study. This work was supported by AMED under Grant Number JP22mk0101250h and 23ak0101199h0001. Additionally, we utilized ChatGPT and Gemini to improve the clarity and readability of the English language in this manuscript. The authors reviewed and revised the output and take full responsibility for the content of the publication.

## Appendix Supplementary Information

To determine the relationships between two cell names (entities) appearing in the text, we established the following annotation criteria.

### 1. General Principles

#### 1.1 Entity Eligibility and Basic Rules

- **Suitability of Entities:**
  ⮚ Instances where the cell name serves as a modifier for a non-cellular noun (e.g., “**[Mast cell]** colony-forming unit”) are considered invalid.
  ⮚ Phrases like “CD8 positive **[T cell]**” are acceptable.
  ⮚ Phrases connecting distinct cells, such as “B and **[T cells]**,” are counted as two independent entities.
  ⮚ Phrases grouping subtypes under a single head noun, such as “CD8+ and CD4+ **[T cells]**,” are treated as a single cell entity.
- **Definition of Directness and Mediation:**
  ⮚ **Substance Mediation:** Cases where Cell A acts on Cell B via a substance secreted by Cell A or via receptors are considered **Direct Relationships**.
  ⮚ **Indirect Mediation:** Any mediation **other than the direct mechanisms defined above** (e.g., involving a third cell or intervening events) is considered indirect and annotated as **No Relation**.
- **Marker Expression:**
  ⮚ Statements merely indicating marker possession (e.g., “positive for cell A marker”) are not considered relationships.
  ⮚ However, if interaction is implied via the marker (e.g., “binding via marker”), it is considered a relationship.

#### 1.2 Contextual Judgment

- **Presence and Strength of Relationship:**
  ⮚ Negative expressions suggesting a lack of relationship (e.g., “poorly,” “few,” “less”) are annotated as **No Relation**.
  ⮚ Positive or suggestive expressions (e.g., “a few,” “suggested”) are annotated as **Relation Exists.**
- **Changes and Effects:**
  ⮚ Contexts describing changes in count, activity, or binding (e.g., “decreases,” “increases”) due to the action of one cell are considered relationships.
- **Semantic Interpretation:**
  ⮚ Even if the grammatical subject or object is not the cell itself, a relationship is determined if the meaning involves the cell’s function or behavior.
  ⮚ **Example:** “Increased binding of T cells to **[B cells]** increases the apoptosis rate of **[macrophages]**.” (Treated as a relationship between B cells and Macrophages).

### 2. Classification Categories

The relationships between extracted cell pairs are classified into the following categories:

#### 2.1 Bidirectional Effect

Refers to undirected or bidirectional physical/functional contact.

- **Definition:** Binding, recognition, or cross-regulation.
- **Examples:**
  ⮚ Binding (where direction is unspecified).
  ⮚ Recognition leading to a change.
  ⮚ Bidirectional regulation (cross-regulation).
  ⮚ Action of B in the presence of A (if directionality is emphasized, “Regulation” may be used; otherwise, “Interaction”).

#### 2.2 Unidirectional Effect

Refers to functional influence with clear directionality.

- **Definition:** One cell causes a change in state, activation, or inhibition of another cell.
- **Examples:**
  ⮚ Expressions indicating direction, such as “effect on.”
  ⮚ “A mediated by B” (Interpreted as B controlling A).

#### 2.3 Change

Refers to the process where one cell changes into another.

- **Definition:** Differentiation or transformation from Cell A to Cell A’ (or B).
- **Examples:** Differentiation from progenitor cells to mature cells.

#### 2.4 No Relation

Refers to cases that do not fit the above categories or where a relationship is explicitly denied.

- Definition:
  ⮚ Simple co-occurrence without described interaction.
  ⮚ Taxonomical relationships like “A is a B”.
  ⮚ Explicitly indirect relationships mediated by another cell.

#### 2.5 Invalid

Refers to cases unsuitable for judgment in this study.

- **Examples:**
  ⮚ The number of target cells is not two (applicable only to SemMed data).
  ⮚ Descriptions of study purpose or methods.
  ⮚ Explicit descriptions of an in vitro setting.

---

**Supplementary Figure 1.**
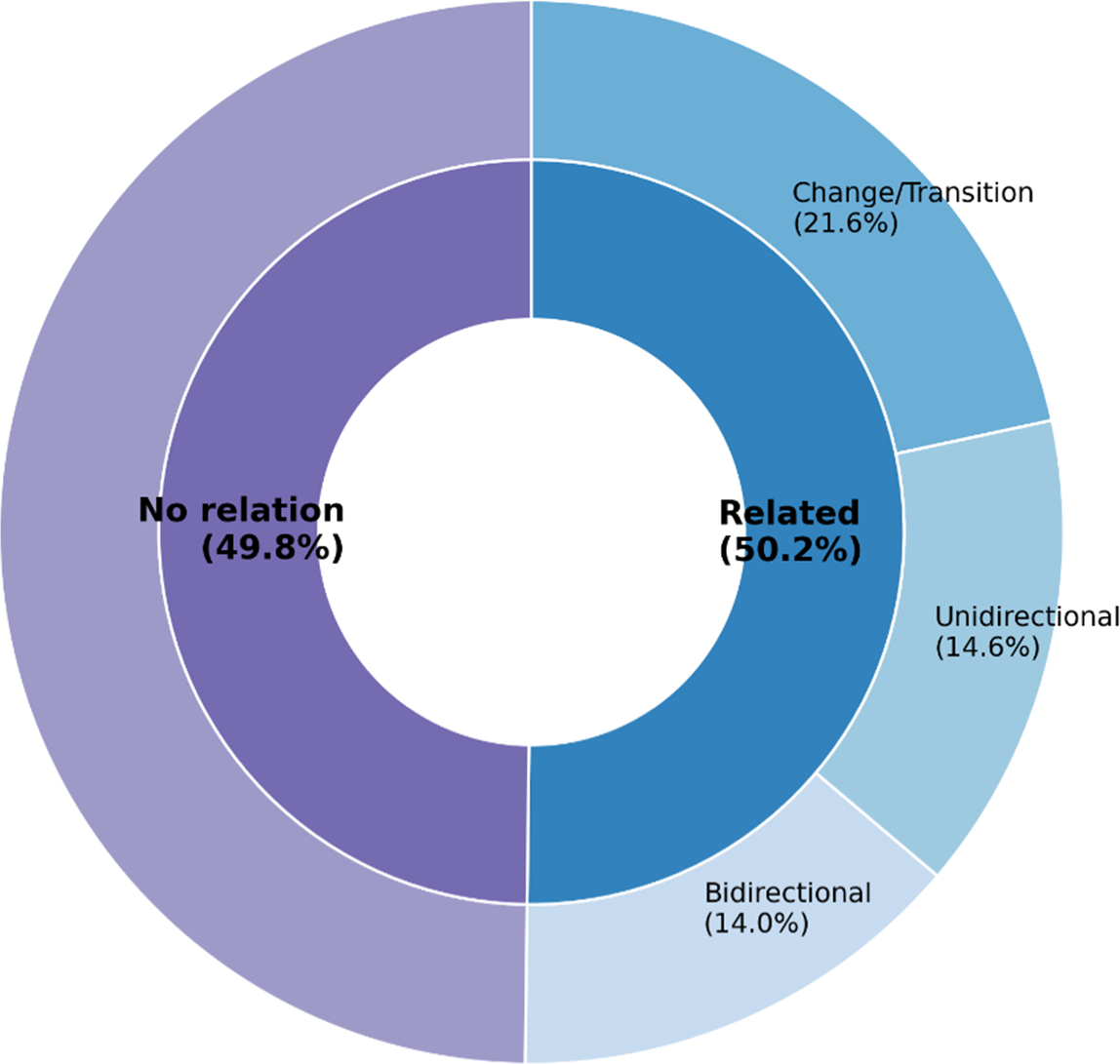
Distribution of relation labels in the SemMed-based corpus. The inner ring represents the proportion of binary classification labels (Related vs. No relation) used in this study. The outer ring displays the breakdown of the positive (“Related”) instances into three detailed interaction types: Change/Transition, Unidirectional, and Bidirectional.

**Supplementary Figure 2.**
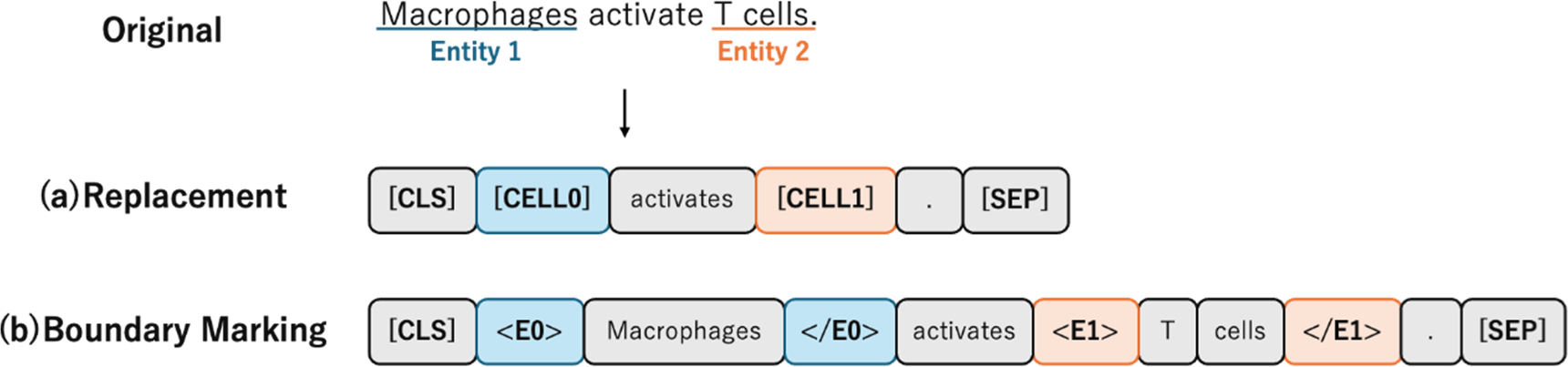
Comparison of Entity Marking Methods. “Original” represents the source sentence. **(a) Replacement:** A method where target cell names are completely replaced with abstract special tokens (e.g., [CELL0], [CELL1]), thereby removing entity-specific lexical information from the input. **(b) Boundary Marking:** A method where boundary tags (e.g., <E0>,</E0>) are inserted surrounding the cell names. This approach explicitly indicates entity positions while preserving the original lexical information.

**Supplementary Figure 3.**
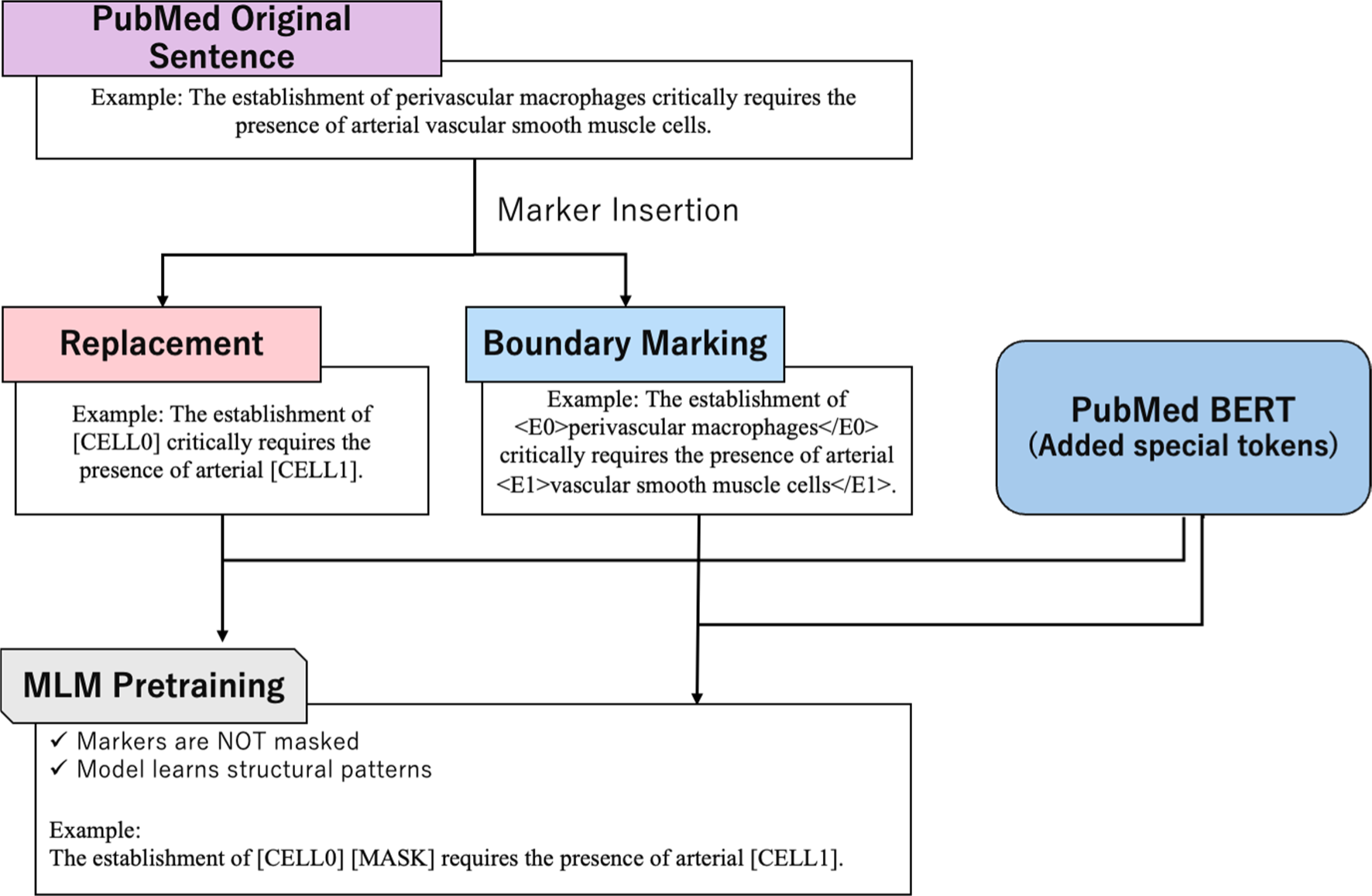
Overview of the Continued Pre-training (CPT) pipeline. Special tokens are inserted into original PubMed sentences using either the Replacement or Boundary Marking method. Subsequently, Masked Language Modeling (MLM) is performed with a model where these special tokens have been added to the vocabulary. Notably, the inserted special tokens are explicitly excluded from masking targets and remain visible. This design allows the model to be exposed to the structural patterns associated with entity markers during pre-training.

**Supplementary Figure 4.**
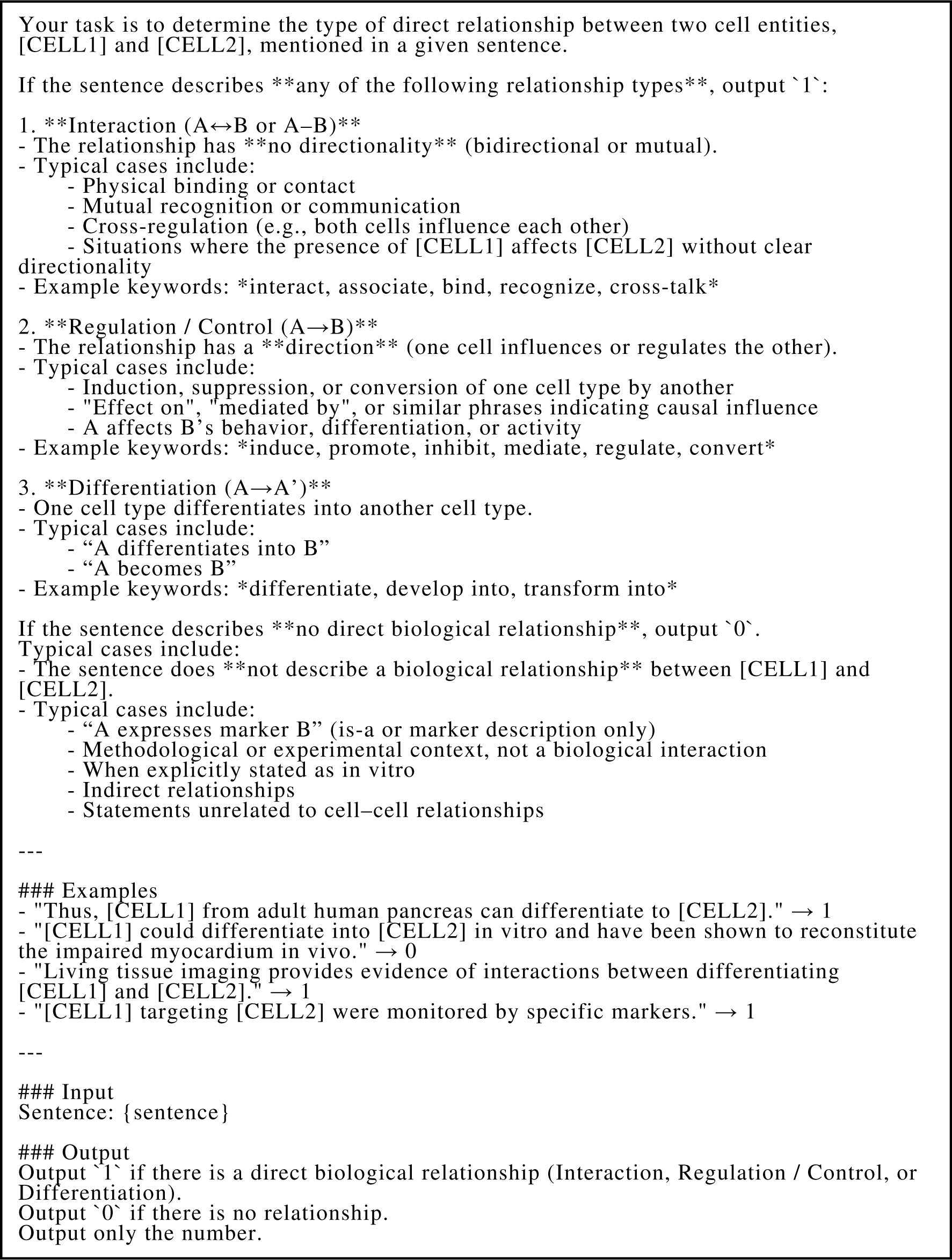
Prompt Design for Large Language Models (LLMs) A standardized prompt was used for all LLMs to perform sentence-level binary classification (Interaction vs. No Relation) given a sentence and two target entities. The prompt instructs the model to perform a binary classification task (Interaction vs. No Relation) given a sentence and two target entities.

**Supplementary Figure 5.**
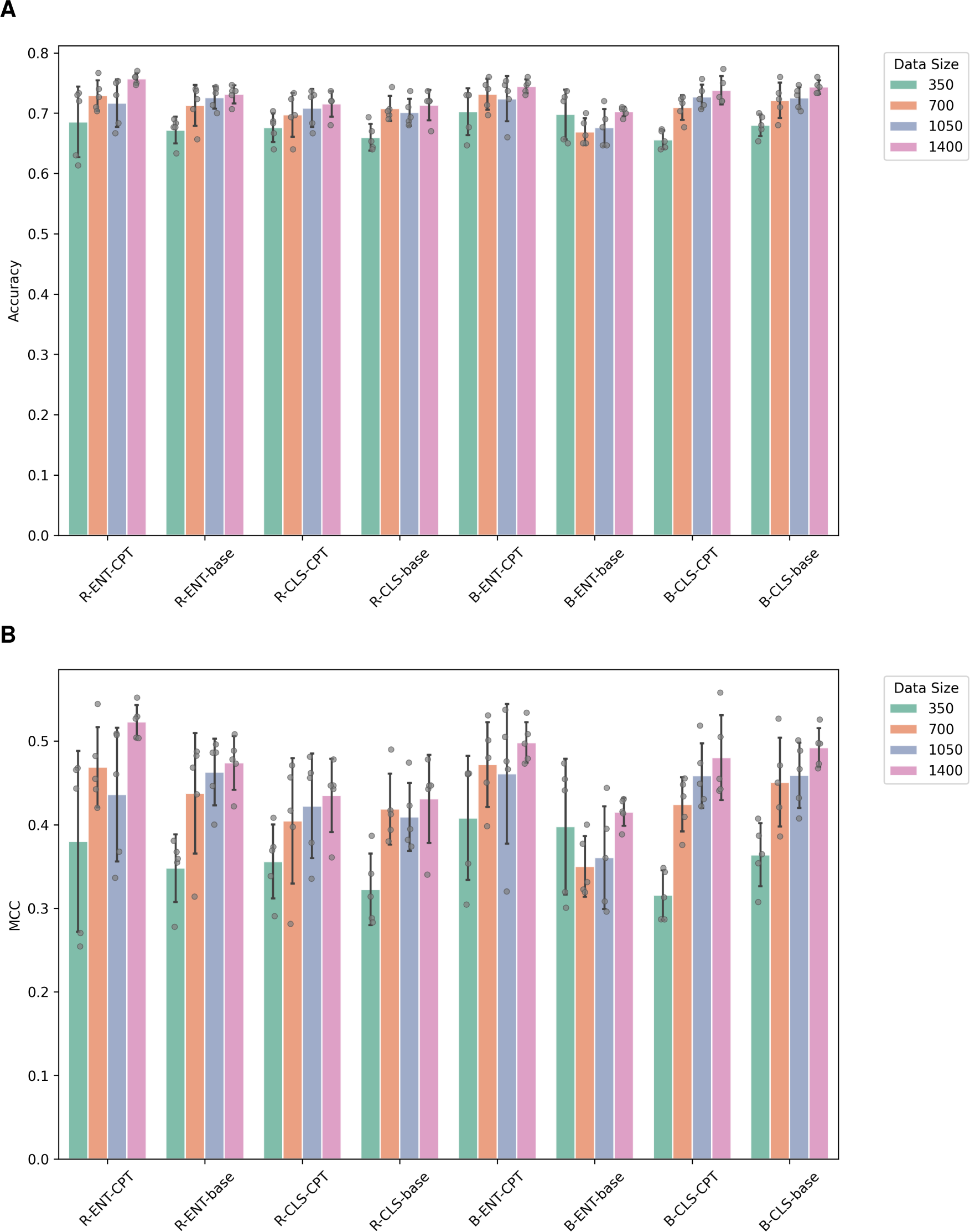
Model performance across different training data sizes. Comparisons of Accuracy (A) and Matthews correlation coefficient (MCC) (B) for different model configurations as a function of training data size. The x-axis represents combinations of classification architecture and pre-training setting (with or without continued pre-training), while the legend indicates the number of training samples used (350, 700, 1050, and 1400).

**Supplementary Figure 6.**
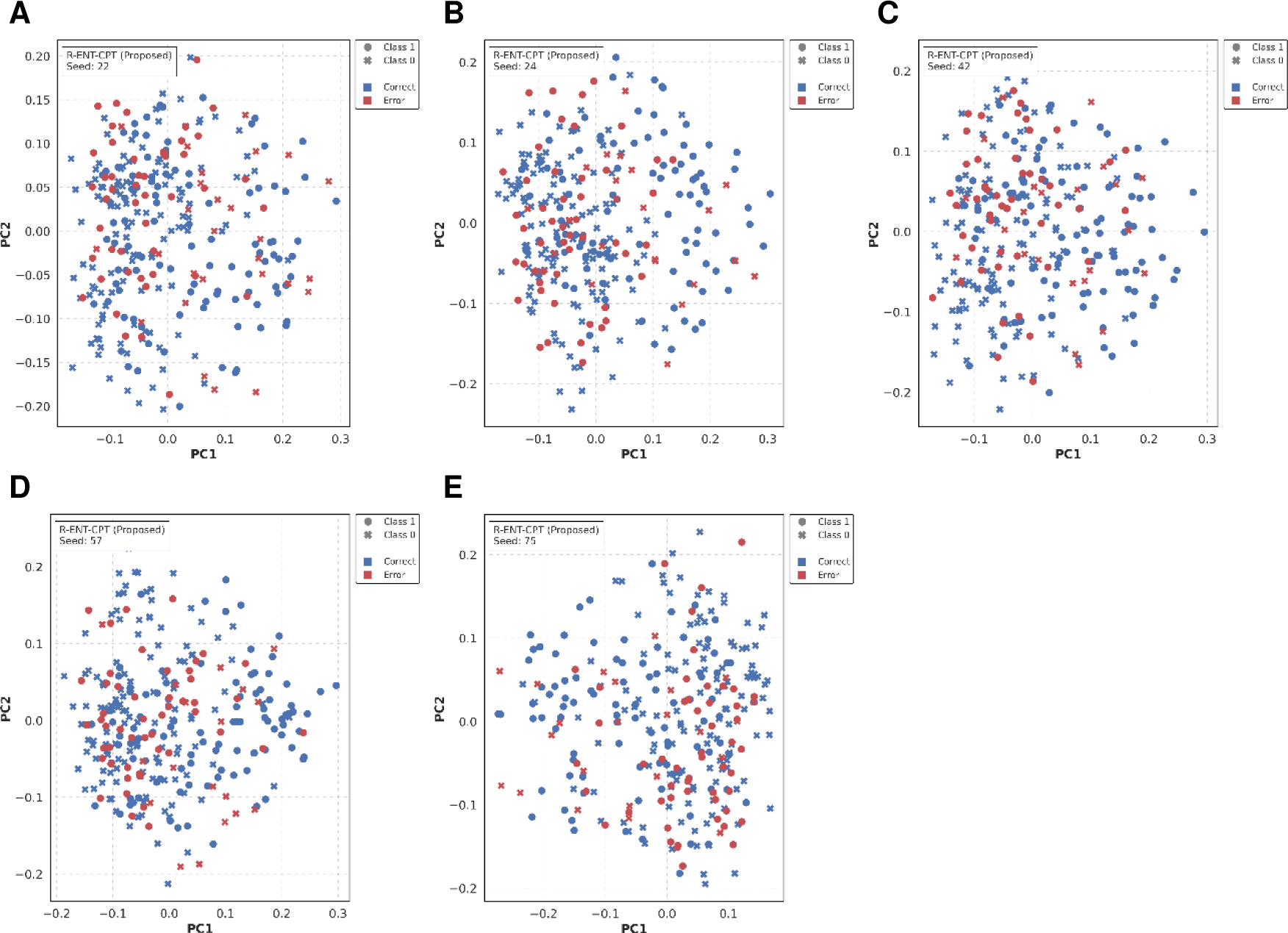
PCA visualization of test embeddings and prediction outcomes. Principal component analysis (PCA) visualization of test data embeddings and prediction results for the proposed model (R-ENT-CPT). Feature vectors are projected into two-dimensional space for visualization purposes. Marker shapes indicate ground truth labels (circle: positive; cross: negative), and colors denote prediction correctness (blue: correct; red: incorrect). Each panel corresponds to a different random seed.

**Supplementary Figure 7.**
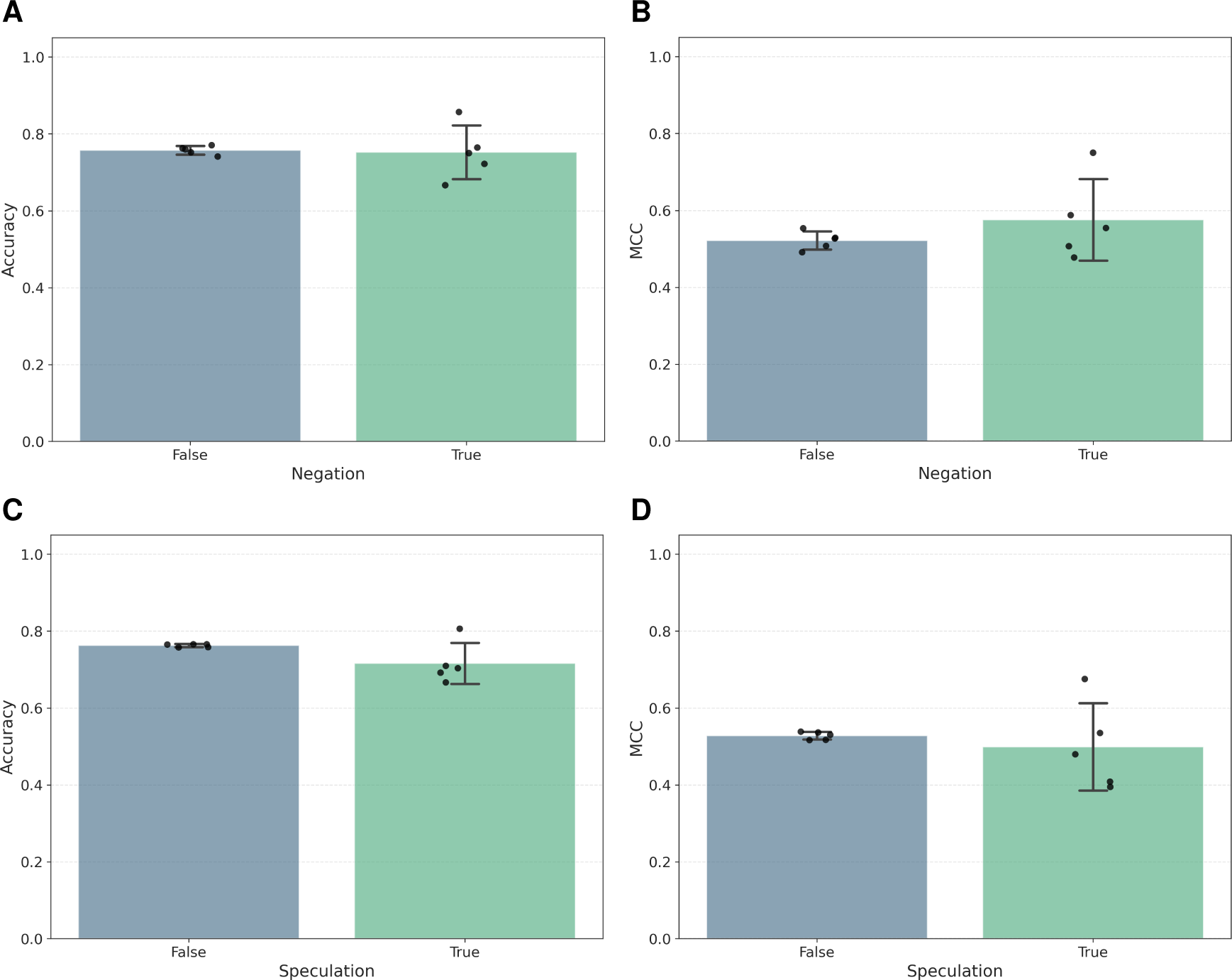
Performance stratified by negation and speculative expressions on the proposed model. Performance of the proposed model (R-ENT-CPT) stratified by the presence of negation (top row) and speculative expressions (bottom row). Distributions of Accuracy (left column) and MCC (right column) are shown. “False” indicates sentences without the corresponding linguistic feature, and “True” indicates sentences containing it.

**Supplementary Figure 8.**
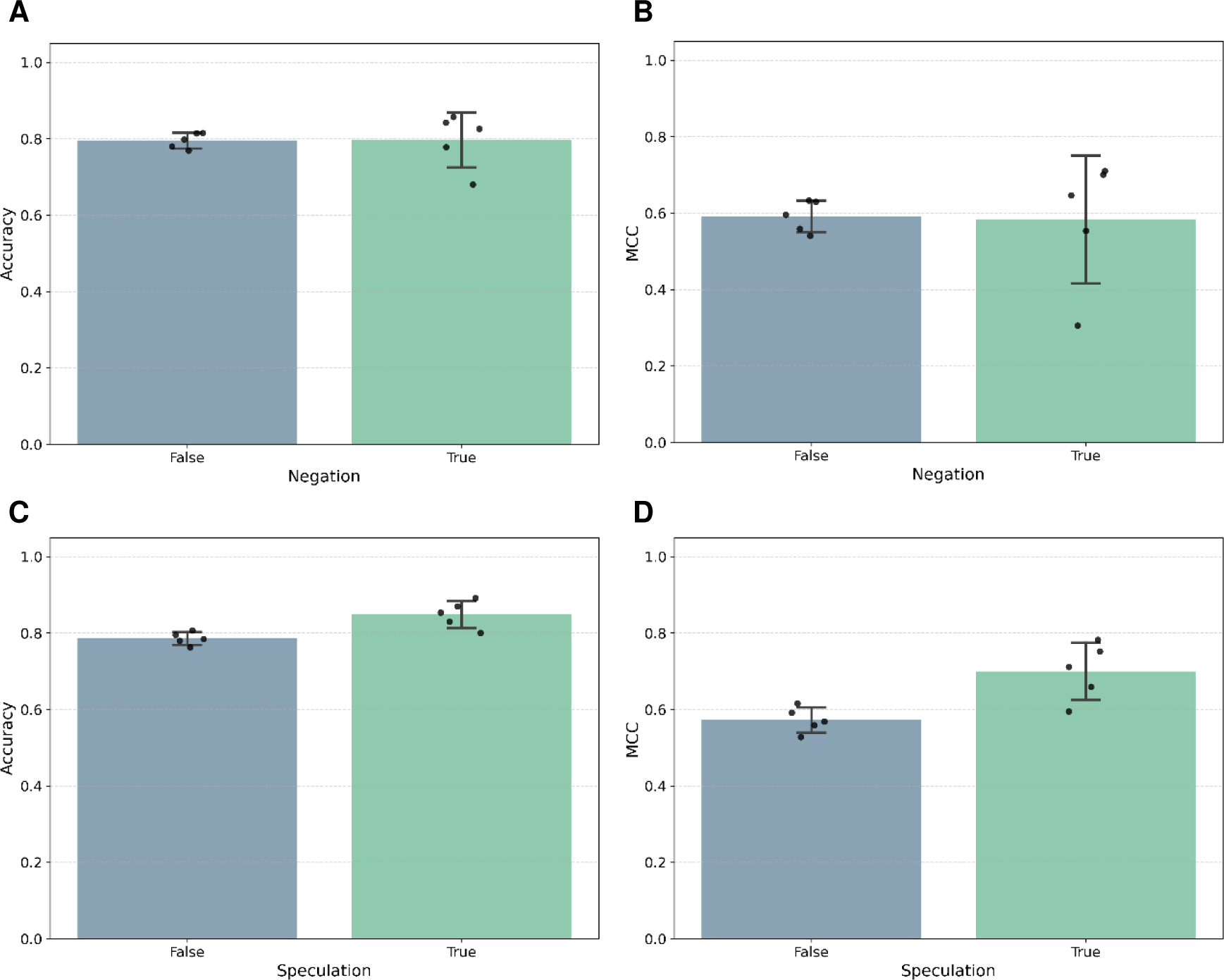
Performance stratified by negation and speculation on GPT-4.1 performance. Performance of GPT-4.1 stratified by the presence of negation (top row) and speculative expressions (bottom row). Accuracy (left column) and Matthews correlation coefficient (MCC) (right column) are shown for sentences with (“True”) and without (“False”) the corresponding linguistic feature.

**Supplementary Figure 9.**
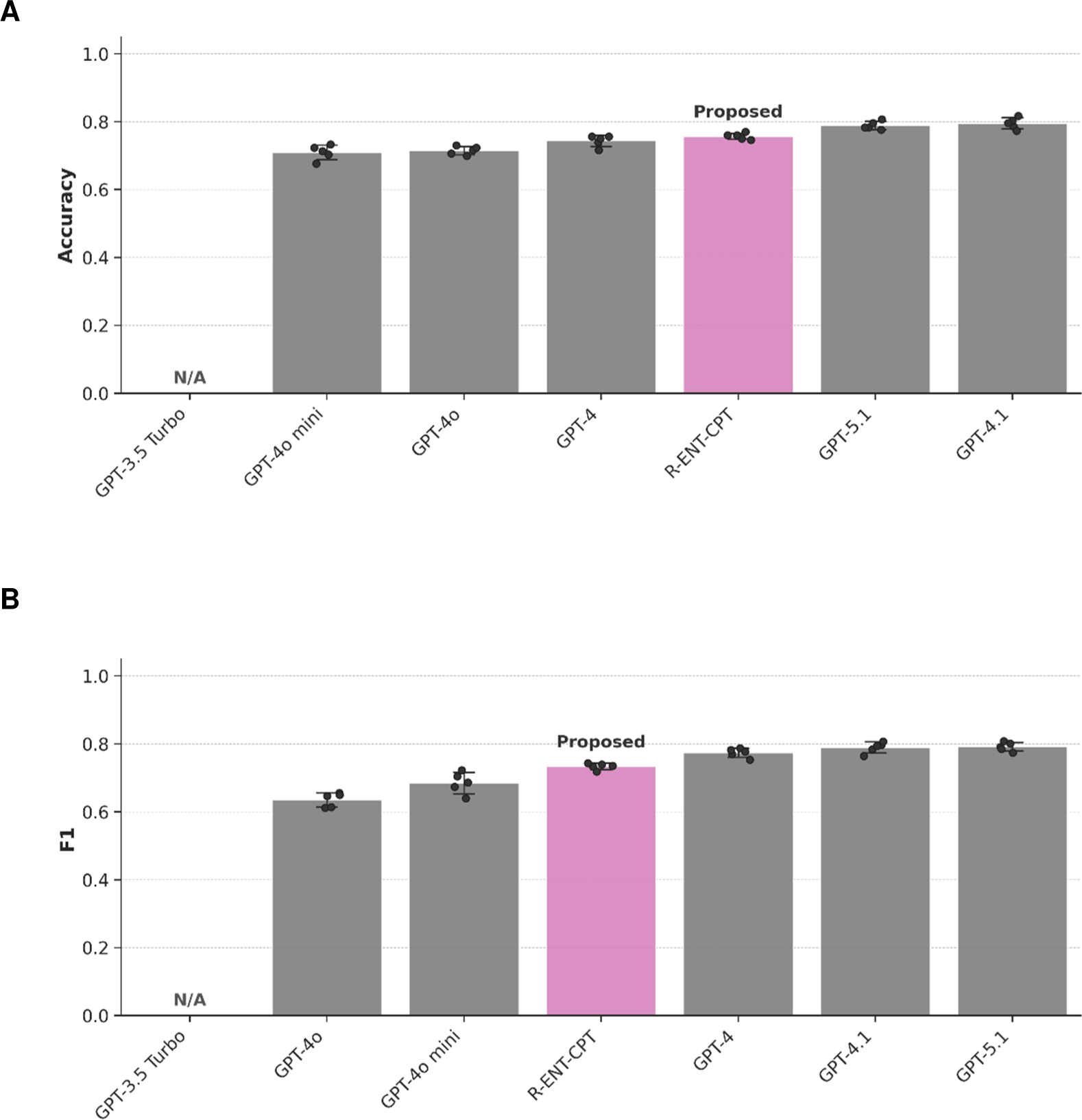
Performance comparison with Large Language Models (LLMs). Evaluation results of the proposed model (R-ENT-CPT) and various OpenAI GPT models on the PubMed test set (OOD data). The panels display (A) Accuracy and (B) F1-score. The pink bar represents the proposed model, while grey bars represent general-purpose LLMs accessed via the OpenAI API. Error bars indicate the standard deviation across multiple runs. Note that GPT-3.5 Turbo is marked as “N/A” as it failed to generate valid outputs following the prompt instructions. This figure provides additional performance metrics corresponding to the comparison shown in **Figure 4**.

**Supplementary Table 1.** Summary of the datasets constructed in this study. The table lists the size (number of sentences), data source, experimental purpose, and text characteristics for each corpus. **CPT:** Continued Pre-training. **ID**: In-Distribution. **OOD:** Out-of-Distribution.

| Corpus | Size | Source | Purpose | Characteristics |
| --- | --- | --- | --- | --- |
| CPT Corpus | 2,121,113 | PubMed | CPT | Marker insertion, large scale |
| SemMed RE Train Corpus | 1,400 | SemMed | Fine-tuning | Structured text, simple syntax |
| SemMed RE Test Corpus | 160 | SemMed | ID Test | Structured text, simple syntax |
| PubMed RE Test Corpus | 300 | PubMed | OOD Test | Natural text, complex syntax |

**Supplementary Table 2.**
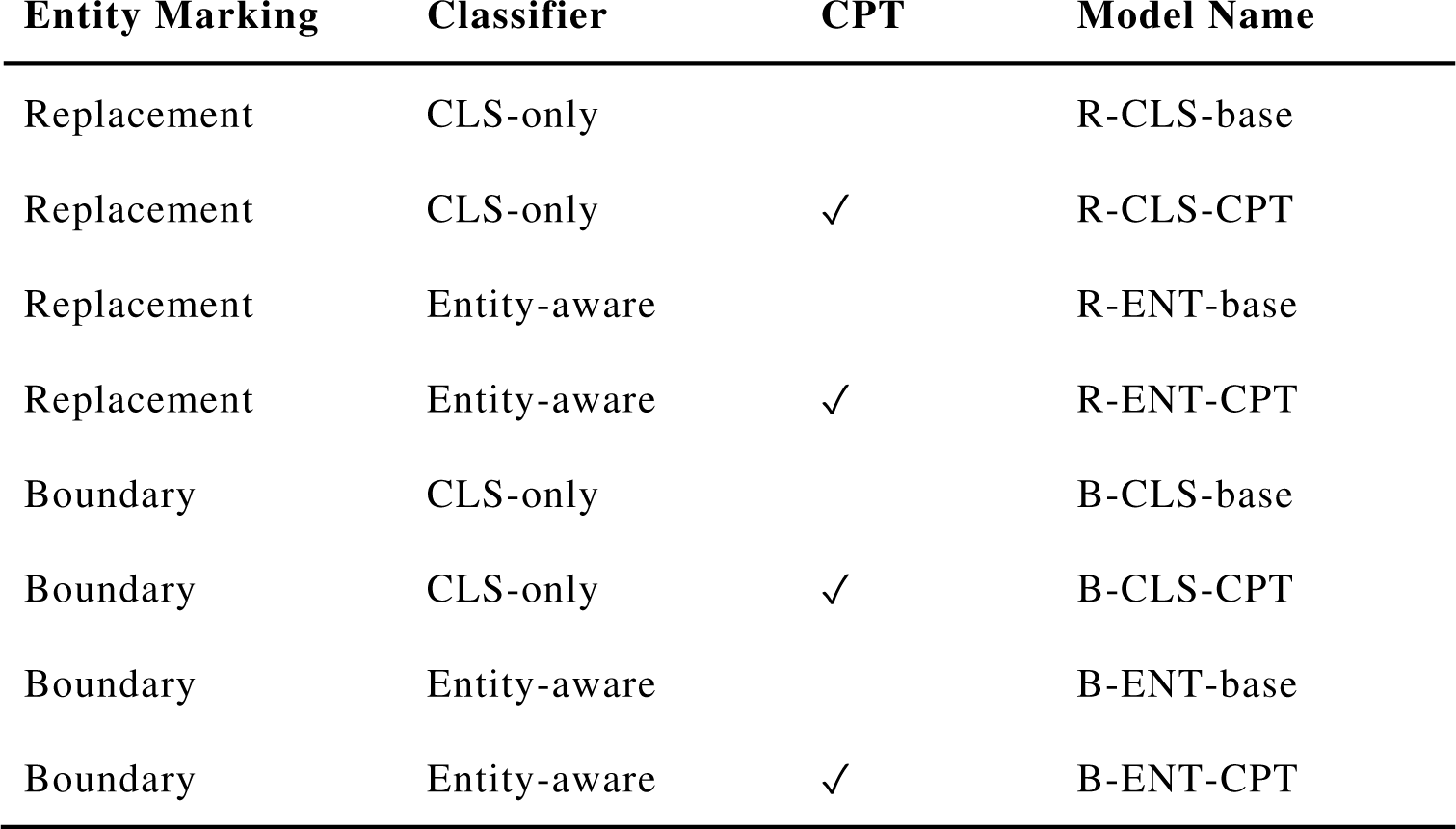
List of model configurations compared in this study. The table lists the combinations of entity marking method, classifier architecture, and the presence or absence of continued pre-training (CPT). The symbol ✓ indicates that CPT was applied. “Model Name” denotes the abbreviations used throughout the experimental results, following the naming convention [Entity Marking]–[Classifier]–[Pre-training].

| Entity Marking | Classifier | CPT | Model Name |
| --- | --- | --- | --- |
| Replacement | CLS-only |  | R-CLS-base |
| Replacement | CLS-only | ✓ | R-CLS-CPT |
| Replacement | Entity-aware |  | R-ENT-base |
| Replacement | Entity-aware | ✓ | R-ENT-CPT |
| Boundary | CLS-only |  | B-CLS-base |
| Boundary | CLS-only | ✓ | B-CLS-CPT |
| Boundary | Entity-aware |  | B-ENT-base |
| Boundary | Entity-aware | ✓ | B-ENT-CPT |

**Supplementary Table 3.** Performance stratified by entity distance. Performance is reported for three entity distance bins (Short: 0–5 tokens, Medium: 6–15 tokens, Long: 16 or more tokens). Values represent mean ± standard deviation for Accuracy and Matthews Correlation Coefficient (MCC).

|  | Proposed |  | GPT-4.1 |  |
| --- | --- | --- | --- | --- |
|  | Accuracy | MCC | Accuracy | MCC |
| Short (0-5) | 0.736±0.022 | 0.482±0.047 | 0.823±0.016 | 0.647±0.028 |
| Medium (6-15) | 0.736±0.028 | 0.510±0.055 | 0.778±0.029 | 0.558±0.060 |
| Long (16+) | 0.796±0.035 | 0.596±0.066 | 0.788±0.053 | 0.574±0.108 |

**Supplementary Table 4.** Performance stratified by sentence length. Performance is stratified by the total number of tokens in the sentence (Short: ≤30 tokens, Medium: 31–45 tokens, Long: 46 or more tokens). Values represent mean ± standard deviation for Accuracy and Matthews Correlation Coefficient (MCC).

|  | Proposed |  | GPT-4.1 |  |
| --- | --- | --- | --- | --- |
|  | Accuracy | MCC | Accuracy | MCC |
| Short (0-5) | 0.743±0.023 | 0.493±0.066 | 0.801±0.027 | 0.597±0.054 |
| Medium (6-15) | 0.775±0.037 | 0.560±0.089 | 0.786±0.045 | 0.572±0.092 |
| Long (16+) | 0.753±0.034 | 0.497±0.065 | 0.798±0.027 | 0.590±0.059 |

**Supplementary Table 5.** Performance comparison with Large Language Models (LLMs). The table reports the performance of the proposed task-specific model (R-ENT-CPT) and several general-purpose LLMs on the out-of-distribution (OOD) PubMed test set. Accuracy, Matthews Correlation Coefficient (MCC), and F1-score are reported. LLMs were evaluated using prompt-based inference without task-specific fine-tuning and are included as reference baselines.

| Model | Accuracy | MCC | F1 |
| --- | --- | --- | --- |
| GPT-4o mini | 0.710±0.022 | 0.425±0.041 | 0.685±0.032 |
| GPT-4o | 0.715±0.012 | 0.480±0.029 | 0.636±0.021 |
| GPT-4 | 0.744±0.017 | 0.506±0.032 | 0.774±0.013 |
| R-ENT-CPT | 0.757±0.009 | 0.523±0.020 | 0.734±0.009 |
| GPT-5.1 | 0.789±0.012 | 0.579±0.024 | **0.792±0.013** |
| GPT-4.1 | **0.795±0.016** | **0.592±0.033** | 0.789±0.016 |

